# Divergent mutational processes distinguish hypoxic and normoxic tumours

**DOI:** 10.1101/531996

**Authors:** Vinayak Bhandari, Constance H. Li, Robert G. Bristow, Paul C. Boutros, on behalf of the PCAWG Network

## Abstract

Many primary tumours have low levels of molecular oxygen (hypoxia). Hypoxic tumours are more likely to metastasize to distant sites and respond poorly to multiple therapies. Surprisingly, then, the pan-cancer molecular hallmarks of tumour hypoxia remain poorly understood, with limited understanding of its associations with specific mutational processes, non-coding driver genes and evolutionary features. To fill this gap, we quantified hypoxia in 1,188 tumours spanning 27 cancer types. We show that elevated hypoxia is associated with increased mutational load across cancers, irrespective of the underlying mutational class. The proportion of mutations attributed to several mutational signatures of unknown aetiology are directly associated with the level of hypoxia, suggesting underlying mutational processes for these signatures. At the gene level, driver mutations in *TP53, MYC* and *PTEN* are enriched in tumours with high hypoxia, and mutations in *PTEN* interact with hypoxia to direct the evolutionary trajectory of tumours. Overall, this work demonstrates that hypoxia plays a critical role in shaping the genomic landscape of cancer.

## Introduction

Approximately half of all solid tumours are characterized by low levels of molecular oxygen (hypoxia)^1–4^. Sub-regions of hypoxia can result from disrupted oxygen supply: irregular and disorganized tumour vasculature can reduce oxygen availability^5^. Hypoxia can also be caused by changes in oxygen demand: altered tumour metabolism^6,7^ can increase intra-cellular demand for oxygen, potentially extending hypoxia signalling to liquid tumours. The adaptation of tumour cells to this imbalance in oxygen supply and demand is associated with poor clinical prognosis in several cancer types, attributed at least in part to hypoxia-associated genomic instability and clonal selection^8–16^.

Previous work has provided insight into the molecular origins and consequences of tumour hypoxia and genomic instability. Dynamic cycling of hypoxia can select for cells with *TP53* mutations and for those that are apoptosis-deficient^17,18^. Indeed mutations in *TP53* occur at a higher frequency in hypoxic primary tumours of at least 9 types^16^. The abundance of proteins involved in homologous recombination (*e.g.* RAD51) and hon-homologous end joining (*e.g.* Ku70) are reduced under hypoxia, and these changes can persist for two days after reoxygenation^19–21^. Genes central to efficient mismatch repair (*e.g. MLH1* and *MSH2*) are also downregulated under hypoxia^22,23^. Further, co-presence of tumour hypoxia and high genomic instability^14,15^, specific cellular morphologies like intraductal and cribriform carcinoma^24^ or specific mutations like loss of *PTEN*^16^, synergistically predict for rapid relapse after definitive local therapy in some tumour types, particularly prostate cancer. These data underscore the relationship between hypoxia and DNA repair defects, and suggest the tumour microenvironment applies a selective pressure leading to the development of specific genomic profiles.

We previously evaluated the exomic and copy-number alteration (CNA) consequences of tumour hypoxia across 19 cancer types^16^. However, the influence of tumour hypoxia on pan-cancer driver alterations, mutational signatures and subclonal architectures remains unclear. To fill this gap, we calculated tumour hypoxia scores for 1,188 tumours with whole-genome and RNA-sequencing sequencing, spanning 27 cancer types. This high-quality harmonized dataset represents a powerful hypothesis-generating mechanism to suggest useful back-translational *in vitro* experiments and better define the hypoxia-associated mutator phenotype. We associated hypoxia with key driver alterations in coding and non-coding regions of the genome, and find hypoxia is associated with specific mutational signatures of unknown aetiology. We illustrate the joint impact of *PTEN* and the tumour microenvironment in influencing the evolutionary trajectory of tumours. Overall, these data highlight the genomic changes through which hypoxia drives aggressive cancers.

## Results

### The pan-cancer landscape of tumour hypoxia

We compiled a cohort of 1,188 tumours from 27 cancer types in the Pan-Cancer Analysis of Whole Genomes (PCAWG) dataset with matched tumour/normal whole-genome sequencing and tumour RNA sequencing data. Whole-genome sequencing^25^ and RNA-sequencing^26^ analyses were systematically carried out by centralized teams with consistent bioinformatics pipelines. Normal samples had a mean whole-genome sequencing coverage of 30 reads per base-pair while coverage for tumour samples had a bimodal distribution with maxima at 38 and 60 reads per base-pair^25^. All samples underwent an extensive and systematic quality assurance process^25^.

We used linear mixed-effect models to associate hypoxia with features of interest across cancers while adjusting for age. Cancer type and sex were further incorporated as random effects in every model, allowing us to consider a different baseline value for the feature of interest for each cancer type and sex^27^. As a measure of effect size, we report 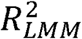 values which reflect the variance explained by the fixed and random factors in each model^28^.

We scored tumour hypoxia in all 1,188 tumours using a trio of mRNA-based hypoxia signatures from Buffa^29^, Winter^30^ and Ragnum^31^ (**Figure 1a**, **Supplementary Figure 1a-b**, **Supplementary Table 1-2**). Hypoxia scores from each of these independent signatures were strongly correlated (ρ = 0.71 – 0.88, all p < 2.2 x 10^-16^, AS89; **Supplementary Figure 1c**) and consistently predicted squamous tumours of the lung (Lung-SCC), cervix (Cervix-SCC) and head (Head-SCC) as the most hypoxic (**Supplementary Figure 1d-e**). Comparatively, chronic lymphocytic leukemias (Lymph-CLL) and thyroid adenocarcinomas (Thy-AdenoCA) were the least hypoxic, consistent with previous^16^ reports (ρ = 0.94, p < 2.2 x 10^-16^, AS89; **Figure 1b**, **Supplementary Figure 1f-h**). Remarkably, subsets of patients from 23/27 cancer types have tumours with elevated hypoxia (hypoxia score > 0) and tumours consistently have elevated hypoxia compared to normal tissues (**Supplementary Figure 2a-c**).

**Figure 1.**
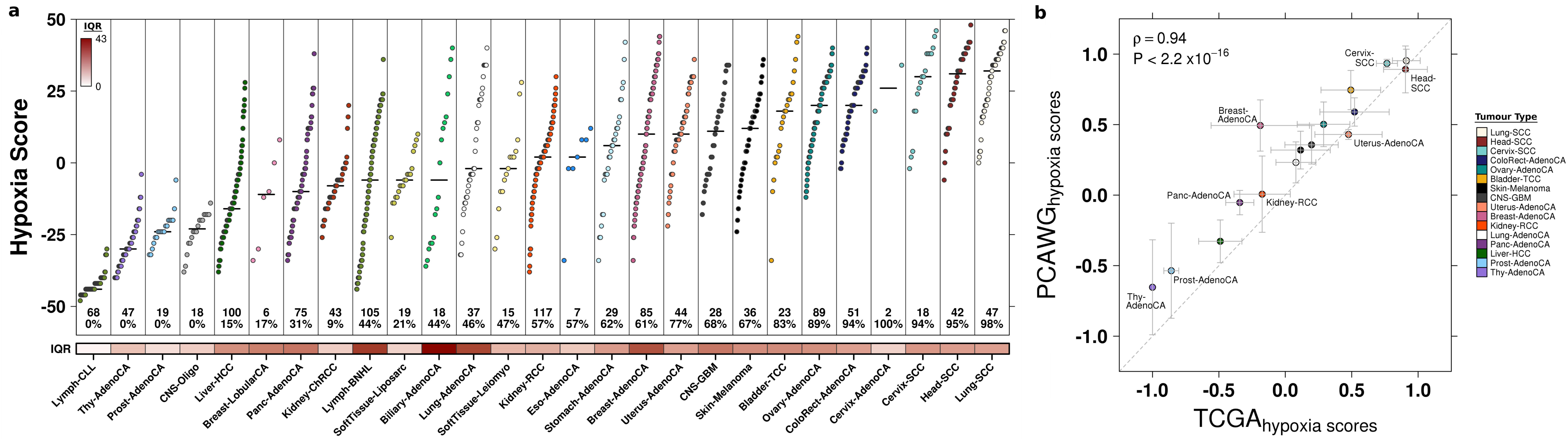
The pan-cancer landscape of tumour hypoxia. We quantified tumour hypoxia in 1,188 tumours spanning 27 different cancer types. **a)** Hypoxia scores for 27 types of cancer, based on the Buffa mRNA abundance signature. Cancer types are sorted by the median hypoxia score (horizontal black line) for each cancer type. Each dot represents one tumour. Sample sizes for each cancer type are listed near the bottom along with the percent of tumours that have elevated hypoxia (hypoxia score > 0). The variability in hypoxia within cancer types was measured by the interquartile range (IQR), shown along the bottom. The IQR was particularly high in biliary adenocarcinoma (IQR = 43.0; Biliary-AdenoCA), mature B-cell lymphomas (IQR = 36.0; Lymph-BNHL), lung adenocarcinoma (IQR = 34.0; Lung-AdenoCA) and breast adenocarcinoma (IQR = 32; Breast-AdenoCA). By contrast, chronic lymphocytic leukemia (IQR = 2.0; Lymph-CLL) and thyroid adenocarcinoma (IQR = 11.0; Thy-AdenoCA) showed less variance in hypoxia score. **b)** Analysis of hypoxia between 16 comparable cancer types in PCAWG and TCGA. Dots represent the mean of the scaled median hypoxia scores from three different mRNA-based hypoxia signatures. Error bars represent the standard deviation of the scaled median hypoxia scores. Overall, the pan-cancer quantification of hypoxia between the PCAWG and TCGA datasets shows strong agreement.

Considering the strong agreement between the Winter, Buffa and Ragnum hypoxia signatures (**Figure 1a**, **Supplementary Figure 1c-d**), we used the Buffa signature for subsequent analyses. We first assessed the degree of inter-tumoural heterogeneity in hypoxia that lies within individual cancer types rather than between them. Over 42% of the variance in hypoxia scores occurs within individual cancer types, highlighting the microenvironmental diversity between tumours arising in a single tissue. This variability in hypoxia score within cancer types was especially elevated in some tumour types, particularly biliary adenocarcinomas (interquartile range, IQR = 43.0; Biliary-AdenoCA), mature B-cell lymphomas (IQR = 36.0; Lymph-BNHL), lung adenocarcinomas (IQR = 34.0; Lung-AdenoCA) and breast adenocarcinomas (IQR = 32.0; Breast-AdenoCA). This was in contrast to chronic lymphocytic leukemias (IQR = 2.0; Lymph-CLL) and prostate adenocarcinomas (IQR = 6.0; Prost-AdenoCA) where little inter-tumoural variability in hypoxia was observed. The variability in hypoxia score was not significantly associated with the median hypoxia score within cancer types (ρ = 0.20, p = 0.30, AS89; **Supplementary Figure 2d**) or with sample size (ρ = 0.22, p = 0.27, AS89; **Supplementary Figure 1e**). Overall, extensive heterogeneity exists in hypoxia levels within and across cancer types.

### The genomic correlates of tumour hypoxia

To determine whether genomic instability arising from specific mutational classes is associated with hypoxia, we looked to identify hypoxia-associated pan-cancer mutational density and summary features^32^. We first considered as a positive control the <u>p</u>ercentage of the <u>g</u>enome with a copy-number <u>a</u>berration (PGA), an engineered feature that is a surrogate for genomic instability and is associated with hypoxia across several tumour types^16^ (**Supplementary Figure 2f**). Indeed, in this diverse pan-cancer cohort, hypoxic tumours have elevated genomic instability while controlling for cancer type, age and sex^27^ (p = 5.01 x 10^-8^, 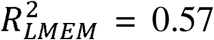, linear mixed-effect model; **Figure 2a**).

**Figure 2.**
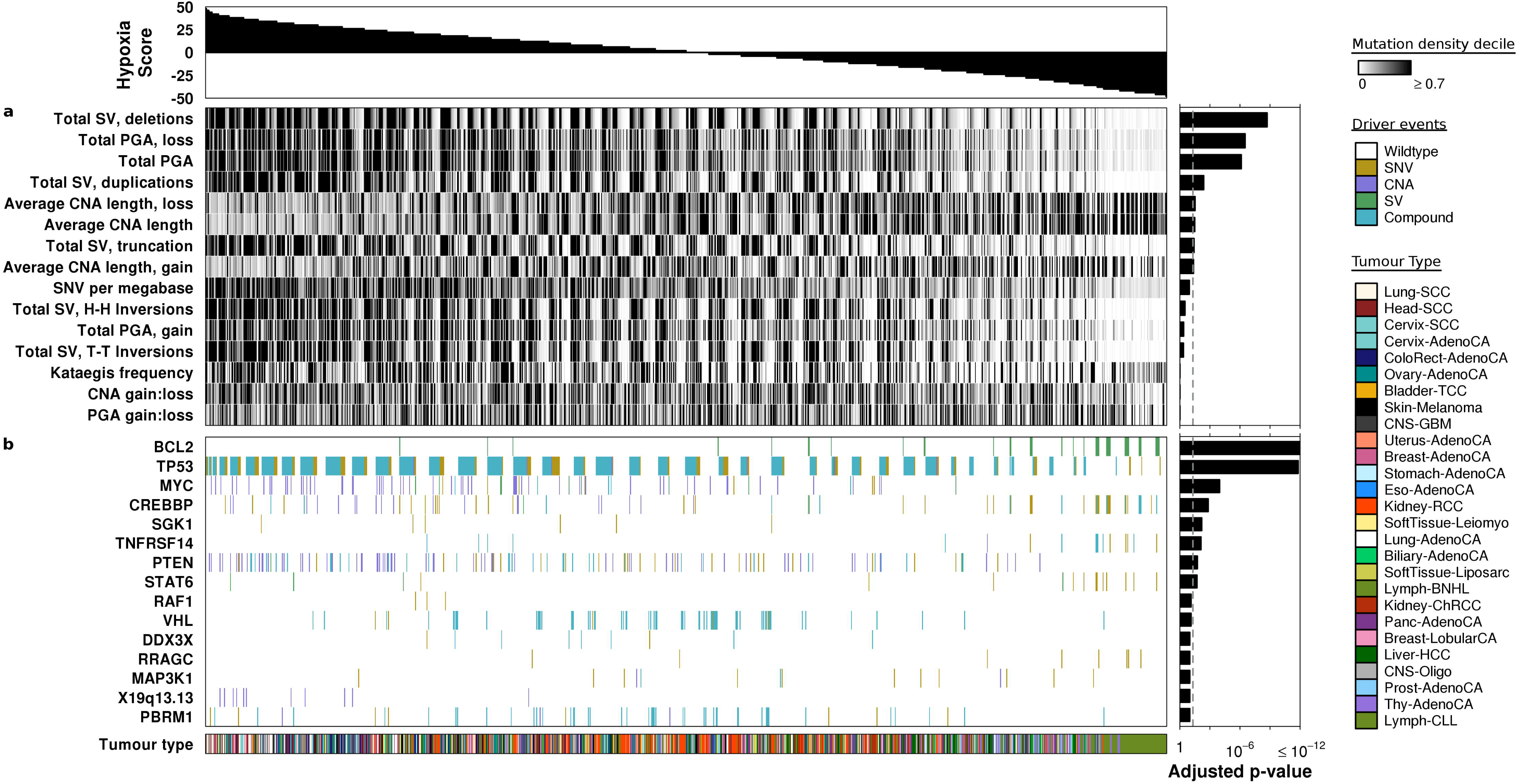
The genomic correlates of tumour hypoxia. We associated tumour hypoxia with mutational density/summary features (**a**) and driver mutations (**b**) across 27 cancer types using linear mixed-effect models. Hypoxia scores for all 1,188 tumours are shown along the top. **a**) Elevated tumour hypoxia was strongly associated with more deletions, elevated PGA, smaller CNAs and a higher number of SNVs per megabase (n = 1,188 independent tumours). Bonferroni-adjusted p-values are shown on the right. **b**) We tested if driver mutations (*e.g.* any of SNV, CNA, SV or a compound event with more than one type of mutation) were associated with hypoxia in 1,096 patients with driver mutation data. Tumours with mutations in *BCL2* showed lower levels of hypoxia while patients with mutations in *TP53* showed remarkably elevated tumour hypoxia. Other driver mutations associated with elevated hypoxia include the oncogene *MYC* and the tumour suppressor *PTEN.* FDR-adjusted p-values are shown along the right. SV, structural variant; PGA, percentage of the genome with a copy-number aberration; CNA, copy-number aberration; SNV, single nucleotide variant; H-H, head-to-head; T-T, tail-to-tail.

We then considered the association of hypoxia scores with 14 other metrics of the mutation density of CNAs, structural variants (SVs) and single nucleotide variants (SNVs) using linear mixed-effect models (**Figure 2a**, **Supplementary Figure 2f**, **Supplementary Table 3-4**). The strongest single correlate of tumour hypoxia was the total number of deletions, where patients with elevated hypoxia had more deletions (p = 1.31 x 10^-10^, 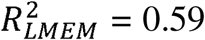, linear mixed-effect model). Elevated numbers of other structural variants such as duplications (p = 2.82 x 10^-4^, 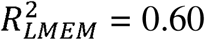, linear mixed-effect model) and truncations (p = 2.58 x 10^-3^, 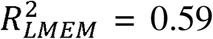, linear mixed-effect model) were also associated with high hypoxia. Other features associated with elevated hypoxia include smaller CNAs (p = 2.22 x 10^-3^, 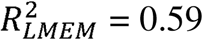, linear mixed-effect model) and more SNVs/Mbp (p = 7.60 x 10^-3^, 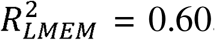, linear mixed-effect model). Overall, hypoxia is associated with increased numbers of most types of somatic mutations.

Considering the strong association of hypoxia with mutational density, we next looked to determine if these were only general effects or selectively affected specific genes or chromosome regions. We leveraged a catalog of 653 driver mutations^33^, with CNA, SV and SNV data available for 1,096 patients. In cases where a patient had multiple mutations in the same gene (*e.g.* a CNA and an SNV) we denoted these as compound events. We again used linear mixed-effect models to associate hypoxia with each driver feature across cancers (**Figure 2b**). Adjusting for cancer type, age and sex, 10 driver events were associated with hypoxia across cancers (FDR < 0.10, linear mixed-effect models; **Supplementary Figure 2f**, **Supplementary Table 5**). Tumours with mutations in *BCL2* (FDR = 7.09 x 10^-15^, 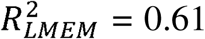, linear mixed-effect model) showed lower levels of hypoxia compared to those without. All alterations of *BCL2* in this cohort were SVs, so it is important to note that this association could not be identified from previous exome-sequencing data. Similarly, mutations in the tumour suppressor *TP53* were associated with elevated hypoxia (FDR = 1.58 x 10^-12^, 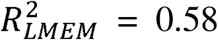, linear mixed-effect model), consistent with previous descriptions of hypoxia-mediated selection of *TP53*-mutated cells^17^ and elevated hypoxia in breast cancer patients with *TP53* mutations^16^. Mutations of the oncogene *MYC* (FDR = 1.09 x 10^-4^, 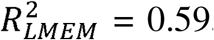, linear mixed-effect model) and tumour suppressor *PTEN* (FDR = 1.80 x 10^-2^, 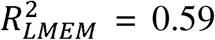, linear mixed-effect model) were also associated with elevated hypoxia. Thus, hypoxia is associated with both broad elevation of mutation density of most types of somatic variation, along with a consistent signature of alterations in oncogenes and tumour suppressors across cancers.

### Hypoxia associated mutational signatures

Previous work has used nonnegative matrix factorization to identify distinct mutational processes in cancer cells from endogenous and exogenous agents^34^. To identify hypoxia-associated mutational processes, we tested if hypoxia score was associated with the proportion of mutations attributed to each mutational signature using linear mixed-effect models. Of the 65 single base substitution (SBS) signatures tested, 9 showed differential activity in hypoxic tumours compared to non-hypoxic ones while controlling for cancer type, age and sex (FDR < 0.10, linear mixed-effect models; **Figure 3a**, **Supplementary Table 6**). Of these, six were more active and three less active in tumours with elevated hypoxia. Since previous work has shown that DNA repair is impaired under hypoxia, it was not surprising to observe that a higher proportion of mutations were attributed to SBS3 in tumours with elevated hypoxia score (FDR = 2.01 x 10^-3^, 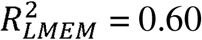, linear mixed-effect model). Further, SBS6 (FDR = 2.01 x 10^-3^, 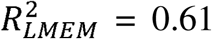, linear mixed-effect model) and SBS21 (FDR = 3.58 x 10^-2^, 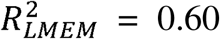, linear mixed-effect model), both related to defective DNA mismatch repair, had a higher proportion of attributed mutations with increasing hypoxia. A lower proportion of mutations were also attributed to SBS1, previously related to the deamination of 5-methylcytosine, with increasing hypoxia (FDR = 2.11 x 10^-7^, 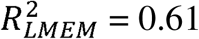, linear mixed-effect model).

**Figure 3.**
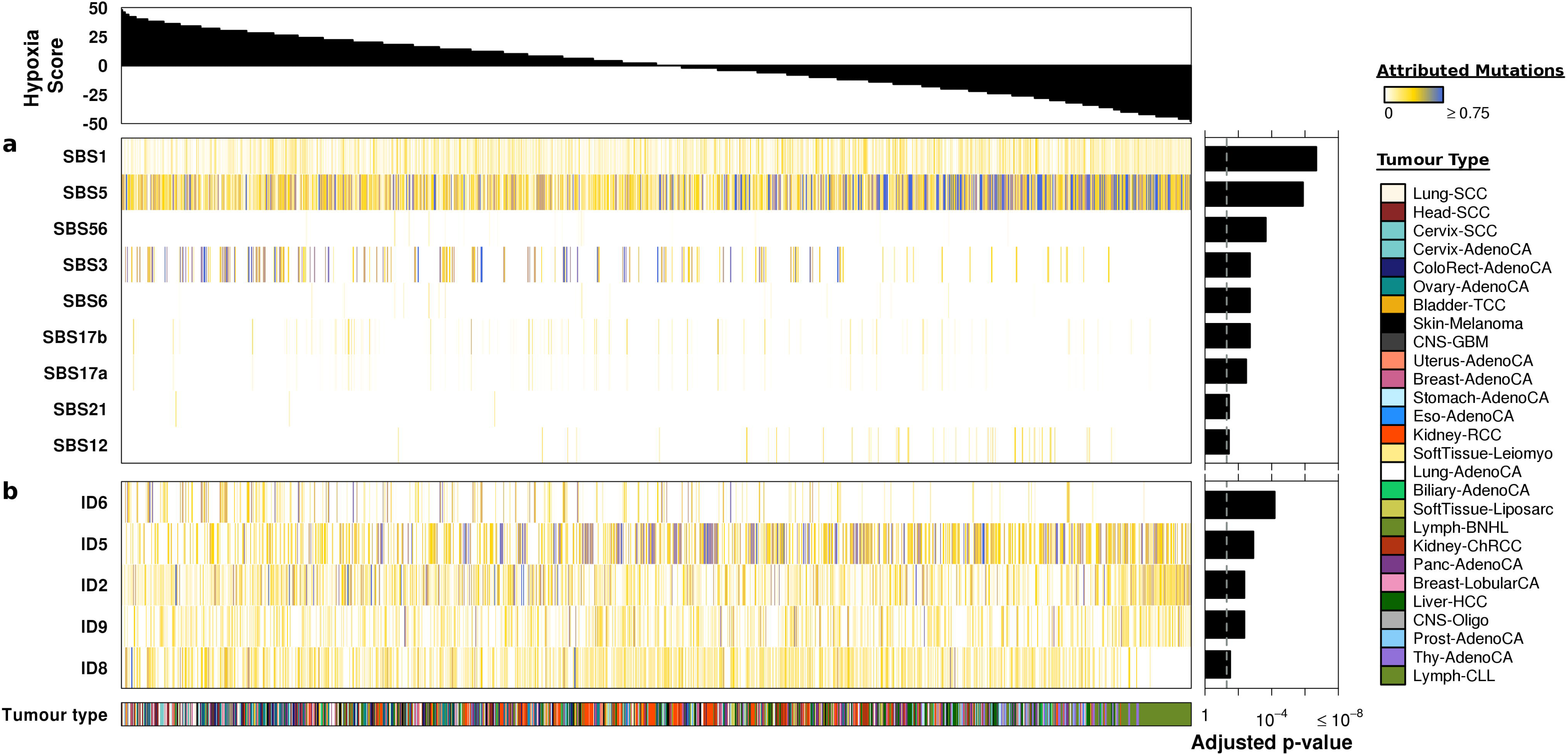
Hypoxia associated mutational signatures. We associated hypoxia with the proportion of mutations attributed to specific mutational signatures using linear mixed-effect models. Hypoxia scores for 1,188 tumours are shown across the top while FDR-adjusted p-values are shown on the right. **a**) Hypoxia was associated with a series of single base substitution signatures with unknown aetiology including SBS5, SBS17a, SBS17b and SBS12. Some of these mutational signatures may reflect hypoxia-dependent mutational processes. Hypoxia was also associated with a lower proportion of attributed mutations to SBS1, which reflects deamination of 5-methylcytosine, and a higher proportion of attributed mutations to SBS3, which is related to deficiencies in DNA double-strand break repair and homologous recombination. **b**) Several signatures of small insertions and deletions were also associated with hypoxia, including ID6 and ID2, which reflect defective homologous recombination and defective DNA mismatch repair, respectively. ID5, ID9 and ID4, all with unknown aetiology, were significantly associated with hypoxia score.

Intriguingly, hypoxia was also associated with a number of SBS signatures with unknown aetiology (**Figure 3b**). The strongest of these was SBS5, where elevated hypoxia was associated with a significantly lower proportion of mutations attributed to the signature (FDR = 1.33 x 10^-6^, 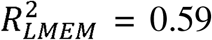 linear mixed-effect model). A significantly lower proportion of mutations were also attributed to SBS12 with increasing hypoxia score (FDR = 3.65 x 10^-2^, 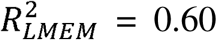, linear mixed-effect model). In contrast, a higher proportion of mutations were attributed to SBS17a (FDR = 3.38 x 10^-3^, 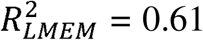, linear mixed-effect model) and SBS17b (FDR = 2.01 x 10^-3^, 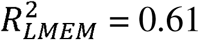, linear mixed-effect model) with increasing hypoxia.

Analysis of small insertion and deletion (ID) signatures illustrated a similar story. Of the 17 ID signatures analyzed, the activity of 5 was associated with tumour hypoxia scores while controlling for cancer type, age and sex (FDR < 0.10, linear mixed-effect models; **Figure 3b**, **Supplementary Table 7**). Of these, 3 were more active in tumours with elevated hypoxia while 2 were less active in them. The defective homologous recombination signature ID6 (FDR = 6.61 x 10^-5^, 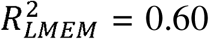, linear mixed-effect model) and defective DNA mismatch repair signature ID2 (FDR = 4.34 x 10^-3^, 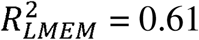, linear mixed-effect model) had a higher proportion of attributed mutations as hypoxia score increased. Several signatures with unknown aetiology were also significantly associated with hypoxia score, including ID5 (FDR = 1.26 x 10^-3^, 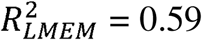, linear mixed-effect model) and ID9 (FDR = 4.34 x 10^-3^, 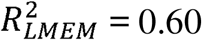, linear mixed-effect model). These data suggest that oxygen levels play a direct or indirect role in the accumulation of specific mutations in cancer cells that are reflected by these signatures.

### The subclonal hallmarks of tumor hypoxia

State-of-the-art methods for subclonal reconstruction rely on whole-genome sequencing data^35^, making the PCAWG dataset an ideal place to understand the evolutionary pressures imposed by hypoxia. Our group and others have shown that some mutations consistently occur early during tumorigenesis while others occur later and that hypoxia is associated with CNAs occurring early in localized prostate cancer^16,36,37^. To explore if this interaction between the tumour microenvironment and mutational landscape exists more broadly in cancer, we assessed if hypoxia was related to the number of clonal or subclonal mutations across 1,188 tumours from 27 cancer types^37^. Clonal mutations are common to all cells in a tumour, while subclonal ones are only present in a subpopulation of cells. We found that elevated hypoxia was significantly associated with an increased number of clonal alterations across cancers (Bonferroni-adjusted p = 4.51 x 10^-3^, 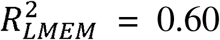, linear mixed-effect model; **Figure 4a**), particularly clonal structural variants (p = 3.19 x 10^-6^, 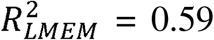, linear mixed-effect model). In contrast, tumour hypoxia was not significantly associated with the number of subclonal alterations (Bonferroni-adjusted p = 0.15, 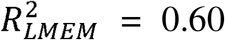, linear mixed-effect model; **Figure 4a**). Further, consistent with previous findings in prostate cancer^16^, hypoxia was not associated with the number of subclones detected (Bonferroni-adjusted p = 0.80, 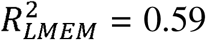, linear mixed-effect model; **Figure 4a**). These data suggest that hypoxia applies a selective pressure on tumours during early tumour development, prior to subclonal diversification.

**Figure 4.**
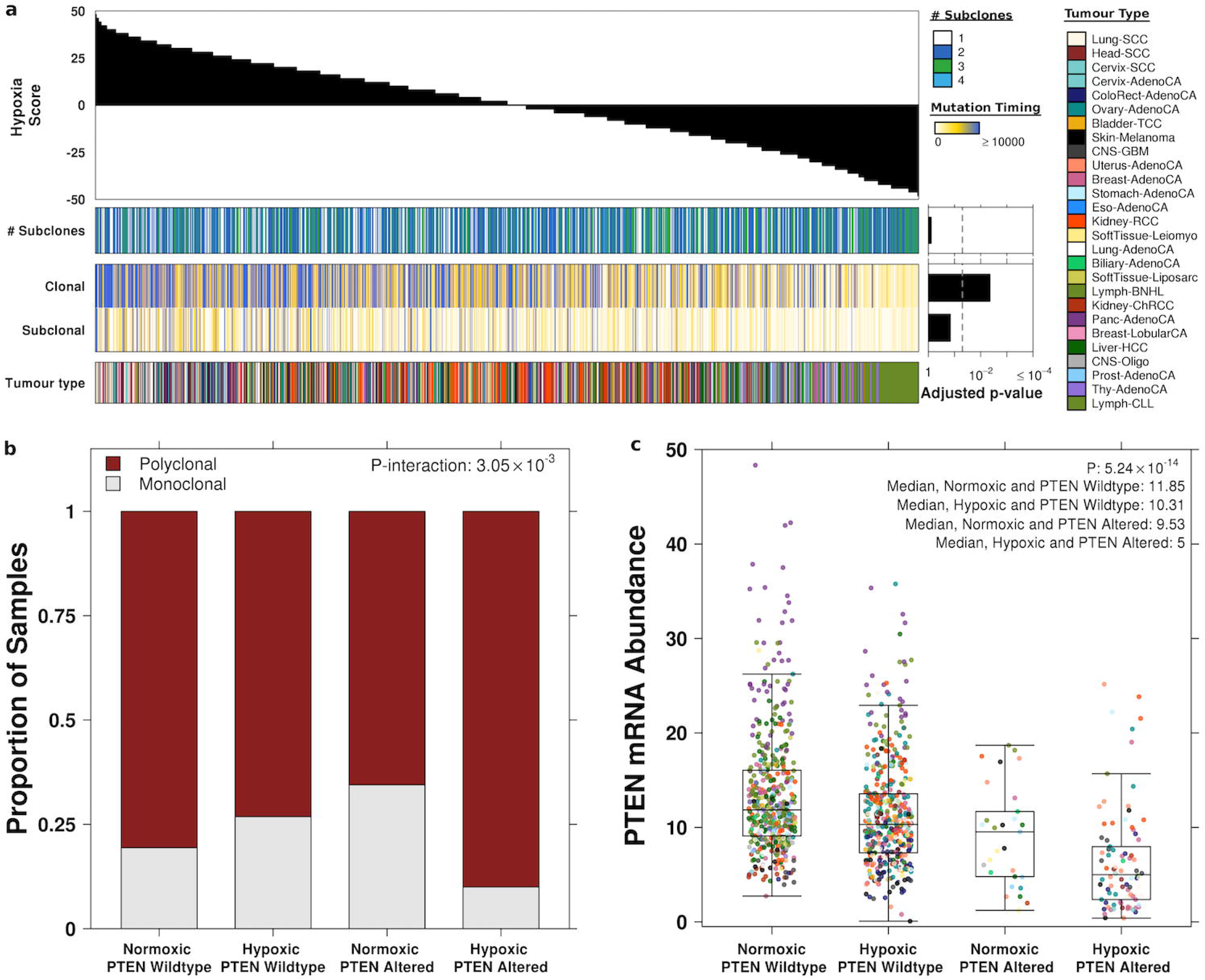
The subclonal hallmarks of tumor hypoxia. We associated tumour hypoxia with features related to the subclonal architecture of 1,188 tumours from 27 cancer types using linear mixed-effect models. **a**) Hypoxia scores are shown along the top while Bonferroni-adjusted p-values are shown on the right. Hypoxia was not associated with the number of subclones in the tumour but elevated hypoxia was associated with a higher number of clonal mutations. **b**) We also observed a significant interaction between hypoxia and altered *PTEN* where tumours with both of these features were particularly likely to be polyclonal. **c**) The mRNA abundance of *PTEN* is modulated by both *PTEN* mutational status and tumour hypoxia. Tumours with altered *PTEN* and elevated hypoxia have the lowest abundance of *PTEN* mRNA.

Next, we assessed if the mutational background of a tumour together with its oxygenation level was linked to its evolutionary trajectory. We previously demonstrated that patients with hypoxic polyclonal prostate tumours with loss of the tumour suppressor *PTEN* tend to have a poor prognosis^16^. Indeed, here we observed a significant interaction between tumor hypoxia and loss of *PTEN* in predicting subclonal architecture (p_interaction_ = 3.05 x 10^-3^, 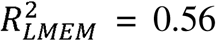, linear mixed-effect model; **Figure 4b**). Specifically, tumours with both of these features tend to have a polyclonal architecture across cancers. The downstream impact of this interaction between the genome and the tumour microenvironment was observed in RNA data: tumours with both altered *PTEN* and elevated hypoxia had the lowest abundance of *PTEN* mRNA (p = 5.24 x 10^-14^, 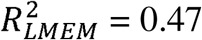, linear mixed-effect model; **Figure 4c**). Thus, the evolutionary trajectory of a tumour may be driven by the presence of a mutation in a specific microenvironmental context (**Figure 5**).

**Figure 5.**
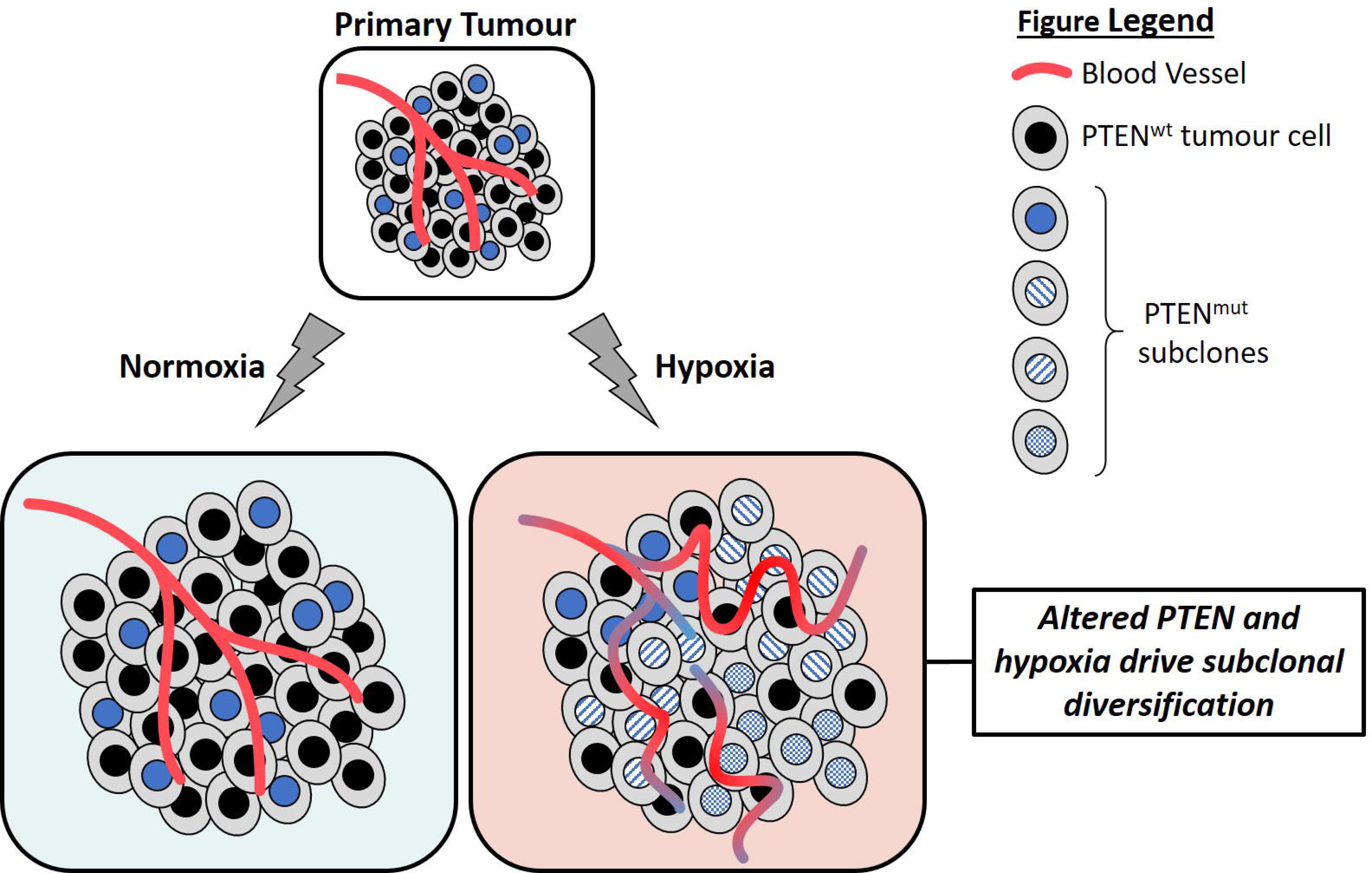
Altered *PTEN* and hypoxia may drive subclonal diversification. Many primary tumours have elevated hypoxia due to increased demand or decreased supply of oxygen. Tumours with elevated hypoxia tend to have altered *PTEN*. Elevated hypoxia and altered *PTEN* may drive subclonal diversification and poor outcomes.

## Discussion

Hypoxia is a feature of many solid and liquid tumours and is associated with aggressive disease. We calculated hypoxia scores for 1,188 tumours from 27 cancer types and showed the vast heterogeneity that exists in this microenvironmental feature within and across cancer types. This reinforces previous pushes for careful patient selection in prospective trials of hypoxia-targeting agents^16^. Further, this work prompts the consideration of basket trials for hypoxia targeting agents to help patients with elevated hypoxia across several cancer types.

For the first time, we characterized the pan-cancer whole-genome correlates of tumour hypoxia. We show the broad influence of the hypoxia associated mutator phenotype: elevated hypoxia is associated with increased mutational load across all mutational classes (*i.e.* CNAs, SVs and SNVs). This supports previous *in vitro* work that demonstrated the contextual synthetic lethality of PARP inhibition in cells with defective DNA repair due to hypoxia^38^. Regarding this co-occurrence of genomic instability and hypoxia, our group^16^ and others^39^ have previously described this metabolic reprogramming as a series of distinct genomic alterations. This is supported by our finding that alterations in *TP53, MYC* and *PTEN* are more common in tumours with elevated hypoxia across cancers. Our study cannot however conclusively say whether hypoxia exerts a selective pressure that enriches for specific genomic alterations or if these genomic changes directly result in hypoxia. Experimental studies of single genes support that both effects may contribute to the associations we describe^17,22,40–42^.

Diving into the mutational processes related to hypoxia, we confirmed that several single base substitution and small indel signatures related to impaired DNA repair were associated with hypoxia. This raises the potential confounder that because hypoxic tumours have more mutations, we have more power to detect related mutational signatures. However, we demonstrated that hypoxia is indeed strongly associated with many mutational signatures with unknown aetiology, particularly SBS5, which is found in nearly all cancer types. Modelling these associations *in vitro* is particularly difficult and these data provide a high confidence measure of the mutational signatures that may be directly or indirectly driven by tumour oxygen levels. It is difficult to disentangle the timing of these events: whether a specific driver mutation gives rise to a specific mutational signature or if these are separate processes. Better mapping of the evolutionary timing of hypoxia will be particularly important in addressing this question and the advent of hypoxia signatures may facilitate future studies in this area.

We observed a significant association between elevated hypoxia and the number of clonal mutations. This supports the idea that hypoxia is an early event in cancer, as we have suggested previously^16^, and other models that link hypoxia to genomic instability and downstream clonal selection^20,41^. Previous work has also demonstrated that patients with allelic loss of *PTEN* and elevated hypoxia rapidly relapse after definitive treatment for localized prostate cancer^16^. Here, we showed that tumours with alterations in *PTEN* and elevated hypoxia are enriched for a polyclonal tumour architecture. This illustrates the joint influence of the tumour mutational landscape and microenvironment in guiding evolutionary trajectories across cancers. Further, these data suggest that increased subclonal diversification may be a novel route *via* which *PTEN* drives aggressive tumour phenotypes, in concert with tumour hypoxia, and this can be better defined with future back-translational *in vitro* experiments. Overall, this work shows that a hypoxic tumour microenvironment is associated with specific mutational processes and distinct somatic mutational profiles, and may direct the subclonal architecture of cancers.

## Methods

### Pan-cancer hypoxia scoring

Hypoxia scores were calculated for all 1,188 tumours with mRNA abundance data using mRNA-abundance based signatures of tumour hypoxia developed previously by Winter *et al.,*^30^ Buffa *et al.*^29^ and Ragnum *et al.*^31^, as described previously^14,16^ (**Supplementary Table 2**). Briefly, patients with the top 50% of mRNA abundance values for each gene in a signature were given a score of +1. Patients with the bottom 50% of mRNA abundance values for that gene were given a score of -1. This was repeated for every gene in the signature to generate a hypoxia score for each patient, and this process was repeated for each of the three signatures used in the study. High scores suggest that the tumour was hypoxic and low scores are indicative of normoxia.

### Hypoxia Score Comparison

To compare hypoxia scores generated by the different signatures, the median hypoxia score was calculated for each of the PCAWG cancer types based on each signature. The median hypoxia scores from each signature were then scaled from +1 to -1 using the plotrix package (v3.7). Scaled median hypoxia values for the PCAWG cancer types were also compared to scaled median hypoxia values from previously published^16^ TCGA data between comparable cancer groups. The groups compared are as follows (PCAWG cancer type TCGA cancer type): Bladder-TCC and BLCA; Breast-AdenoCA and BRCA; Cervix-SCC and CESC; CNS-GBM and GBM; ColoRect-AdenoCA and COADREAD; Head-SCC and HNSC; Kidney-RCC and KIRC; Liver-HCC and LIHC; Lung-AdenoCA and LUAD; Lung-SCC and LUSC; Ovary-AdenoCA and OV; Panc-AdenoCA and PAAD; Prost-AdenoCA and PRAD; Skin-Melanoma and SKCM; Thy-AdenoCA and THCA; Uterus-Adeno and UCEC. Of the 27 cancer types in PCAWG, 11 (Cervix-AdenoCA, Stomach-AdenoCA, Eso-AdenoCA, Breast-LobularCA, SoftTissue-Leiomyo, Lymph-BNHL, SoftTissue-Liposarc, Biliary-AdenoCA, Kidney-ChRCC, CNS-Oligo and Lymph-CLL) did not have hypoxia data from comparable cancers in TCGA and were not used for the comparison^16^. For Spearman’s correlations, p-values were calculated using algorithm AS89.

### Linear Mixed-Effect Models

We used linear mixed-effect models to associate hypoxia with features of interest (*e.g.,* PGA, *TP53* mutational status, etc.) across cancers using the lme4 package (v1.1-17). For each feature of interest we compared a full model (*i.e.,* a model with the feature of interest) to a null model (*i.e.* a model without the feature of interest) using an ANOVA to determine if hypoxia was significantly associated with the feature of interest across cancers:

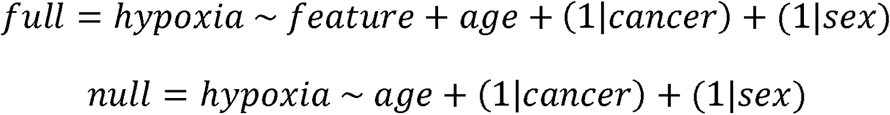

All models were adjusted for patient age. Cancer type and sex were used as random effects in every model. This allows us to consider a different baseline value for the feature of interest for each cancer type and sex^27^. For each model an 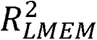 value is reported, reflecting the variance explained by the fixed and random factors^28^.

All model diagnostics were done using the DHARMa package (0.2.0) which uses a simulation-based approach to create standardized residuals^43^. For each model, scaled residuals were generated using the simulateResiduals function. The full model was used as the input for fittedModel parameter and 1,000 simulations were run. For correctly specified models, the scaled residuals were expected to be uniformly distributed and this was verified for each full model. We also compared the standardized residuals to the rank transformed predicted values to assess deviations from uniformity for each full model.

### Mutational Density Analysis

Previously published data for 15 mutational density and summary features were downloaded for 1,188 tumours^32^. We used linear mixed-effect models to associate each feature with hypoxia score across cancers and compared each full model with a null model. Cancer type and sex were used as random effect variables. Tumours belonging to cancer types with fewer than 15 samples were excluded from the analysis. A Bonferroni p-value adjustment was applied to the p-values from linear mixed-effect modeling since fewer than 20 tests were conducted.

### Driver Mutations Analysis

Data for driver mutations was first summarized at the gene level for 1,096 tumours with previously published driver mutation data^33^. For each of the 653 driver features, we summarized if a patient had an SNV, CNA or SV. Some tumours had more than one type of event in a gene (*e.g.* a CNA and an SNV) and these events were classified as compound events. We then used linear mixed-effect models to associate the mutational status of each gene with hypoxia score and compared each full model with a null model. Cancer type and sex were used as random effect variables. The driver mutation analysis did not specifically consider the type of mutation in the gene and only considered if the gene had a mutation or was wildtype. Tumours belonging to cancer types with fewer than 15 samples were excluded from the analysis. An FDR adjustment was applied to the p-values from linear mixed-effect modeling.

### Mutational Signature Analysis

Previously published data for mutations attributed to various specific signatures was downloaded for 1,188 tumours^34^. For each tumour, we calculated the proportion of total mutations attributed to each mutational signature. The proportion of mutations attributed to each signature were calculated by dividing the number of mutations attributed to each signature by the total number of mutations in the tumour. We used linear mixed-effect models to associate the proportion of mutations attributed to each signature with hypoxia score and compared each full model with a null model. Cancer type and sex were used as random effect variables. Tumours belonging to cancer types with fewer than 15 samples were excluded from the analysis. An FDR adjustment was applied to the p-values from linear mixed-effect modeling.

### Subclonality Analysis

Previously reported^37^ subclonal reconstruction data was used to summarize the number of clonal and subclonal mutations in all 1,188 tumours. We used linear mixed-effect models to associate the number of these timed mutations with hypoxia score and compared each full model with a null model. Cancer type and sex were used as random effect variables. Tumours belonging to cancer types with fewer than 15 samples were excluded from the analysis. A Bonferroni adjustment was applied to the p-values from linear mixed-effect modeling since fewer than 20 tests were conducted.

The number of subclones was calculated for all 1,188 tumours based on the number of clusters of cells identified in each sample. A linear mixed-effects model were used to associate the number of subclones with hypoxia score and this model was compared to a null model. Cancer type was used as a random effect. Tumours belonging to cancer types with fewer than 15 samples were excluded from the analysis.

Patients with only one identified cluster of cells were defined as monoclonal and patients with more than one identified cluster of cells were defined as polyclonal^36^. Hypoxia scores were median dichotomized to classify patients as hypoxic or normoxic. To test for an interaction between tumour hypoxia and *PTEN* mutational status in selecting for a particular subclonal architecture, we used linear mixed-effect models together with an ANOVA. A full model was first created where the relationship between the hypoxia scores and *PTEN* mutational status was modelled as an interaction. A reduced model was also created where the relationship between hypoxia scores and *PTEN* mutational status was modelled in an additive manner:

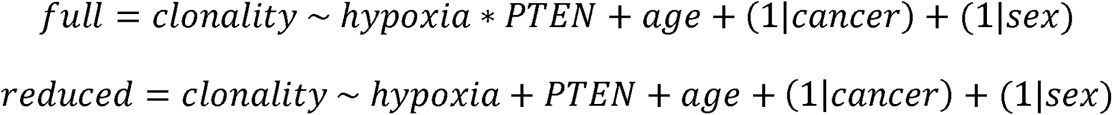

The two models were compared using an ANOVA to test if hypoxia scores significantly interact with *PTEN* mutational status. Tumours belonging to cancer types with fewer than 15 samples were excluded from the analysis. The full model diagnostics were carried out using the DHARMa package, as described above.

All data analysis was performed in the R statistical environment (v3.4.3). Data visualization was performed using the BPG package^44^ (v5.9.1). Figures were compiled using Inkscape (v0.91).

## Authors’ contributions

Bioinformatics and Statistical Analysis: VB

Data Processing: VB, CHL

Data Visualization: VB

Supervised Research: RGB, PCB

Initiated the Project: VB, PCB

Wrote the First Draft of the Manuscript: VB

Approved the Manuscript: All Authors

## Acknowledgements

The authors gratefully thank Dr. Marianne Koritzinsky for insightful suggestions and thank Jenna E. van Leeuwen for visualization support. The authors also thank all members of the Boutros and Bristow labs for helpful suggestions.

## Supplementary Figure Legends

**Supplementary Figure 1 Pan-cancer landscape of hypoxia in primary tumours**

Tumour hypoxia scores for 1,188 tumours from 27 cancer types based on the Winter^30^ (**a**) and Ragnum^31^ (**b**) hypoxia signatures. The black horizontal lines indicate the median hypoxia score within each cancer type. The number of tumours from each cancer type are shown along the bottom. IQR values for each cancer type are also shown along the bottom. **c**) Hypoxia scores based on the three independent signatures were highly correlated for 1,188 tumours from 27 cancer types. **d**) Comparison of median hypoxia scores within cancer types based on the Buffa^29^, Winter^30^ and Ragnum^31^ hypoxia signatures. Median hypoxia values were scaled from +1 to -1. **e**) The mean of the scaled median hypoxia scores for each cancer type are shown by the filled circle Lines show the standard deviation of the scaled median hypoxia score. Squamous tumours of the lung (Lung-SCC), cervix (Cervix-SCC) and head (Head-SCC) are amongst the most hypoxic cancer types. **f-h**) Association of hypoxia scores between 16 comparable types of cancer in PCAWG and TCGA. Dots represent the mean of the scaled median hypoxia scores from the Buffa^29^ (**f**), Winter^30^ (**g**) and Ragnum^31^ (**h**) hypoxia signatures. Pan-cancer calculations of hypoxia between the PCAWG and TCGA datasets is strongly correlated based on independent hypoxia signatures.

**Supplementary Figure 2 Hypoxia in tumour *vs.* normal samples and the genomics of hypoxia across cancers**

Tumour samples consistently have elevated hypoxia compared to normal samples from the same tissue based on the Buffa (**a**), Ragnum (**b**) and Winter (**c**) hypoxia signatures. The IQR values of cancer types were not associated with the median hypoxia score within that cancer type (**d**) or the sample size (**e**). **f**) A partial summary of the knowledge around the genomics of hypoxic tumours. Hypoxia-related associations that have been previously examined within 19 tumour types in TCGA are shown along the left based on data from Bhandari *et al.*^16^. CNAs and SNVs in several genes were found to be associated with elevated hypoxia within tumour types. Hypoxia was also associated with elevated PGA in 10 tumour types. The right side of the figure partially summarizes the analyses carried out in this work. Several of the previous intra-tumour type findings have been extended in this work as pan-cancer features of hypoxia and several novel features, particularly related to structural variants, have been assessed and found to be significantly associated with hypoxia. This is in addition to the novel pan-caner work presented around single-base substitution signatures, indel signatures and tumour subclonality.

## Supplementary Table Legends

**Supplementary Table 1 – PCAWG cancer type codes**

Cancer type codes for the 27 PCAWG cancer types examined in this study and their descriptions.

**Supplementary Table 2 – Hypoxia scores and genomic data for 1,188 tumours**

Data for hypoxia scores, mutational density/summary, driver mutations, mutational signatures and subclonality for 1,188 tumours from 27 cancer types. For driver mutations, 0 represents wildtype, 1 represents an SNV, 2 represents a CNA, 3 represents an SV and 4 represents a compound event (*i.e.,* those with multiple types of alterations). Mutational signature data for single base substitution signatures and insertion and deletion (ID) signatures are provided as the proportion of alterations attributed to the signature.

**Supplementary Table 3 – Mutational density data by deciles**

For each of the 15 mutational features analyzed, the value corresponding to each decile is provided.

**Supplementary Table 4 – Hypoxia associated mutational density features** Results from linear mixed-effect models associating hypoxia with mutational density features while controlling for cancer type, age and sex.

**Supplementary Table 5 – Hypoxia associated driver mutations**

Results from linear mixed-effect models associating hypoxia with driver mutations while controlling for cancer type, age and sex.

**Supplementary Table 6 – Hypoxia associated single base substitution signatures**

Results from linear mixed-effect models associating hypoxia with the proportion of mutations attributed to single base substitution signatures while controlling for cancer type, age and sex.

**Supplementary Table 7 – Hypoxia associated small insertion and deletion signatures**

Results from linear mixed-effect models associating hypoxia with the proportion of mutations attributed to small insertion and deletion signatures while controlling for cancer type, age and sex.

