## Supplementary figures and images for "Divergent mutational processes distinguish hypoxic and normoxic tumours"

### Supplementary Figure 1

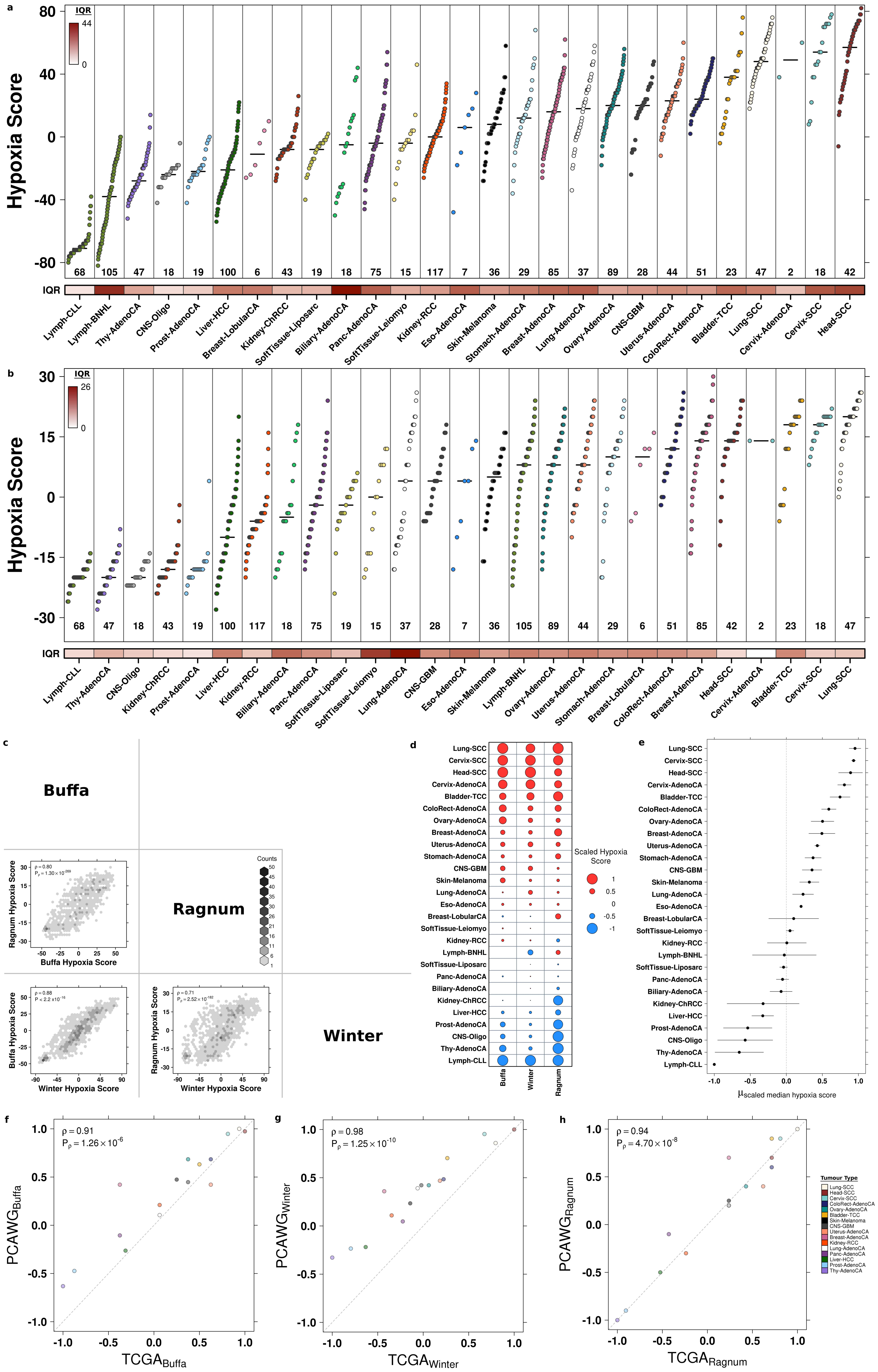

### Supplementary Figure 2

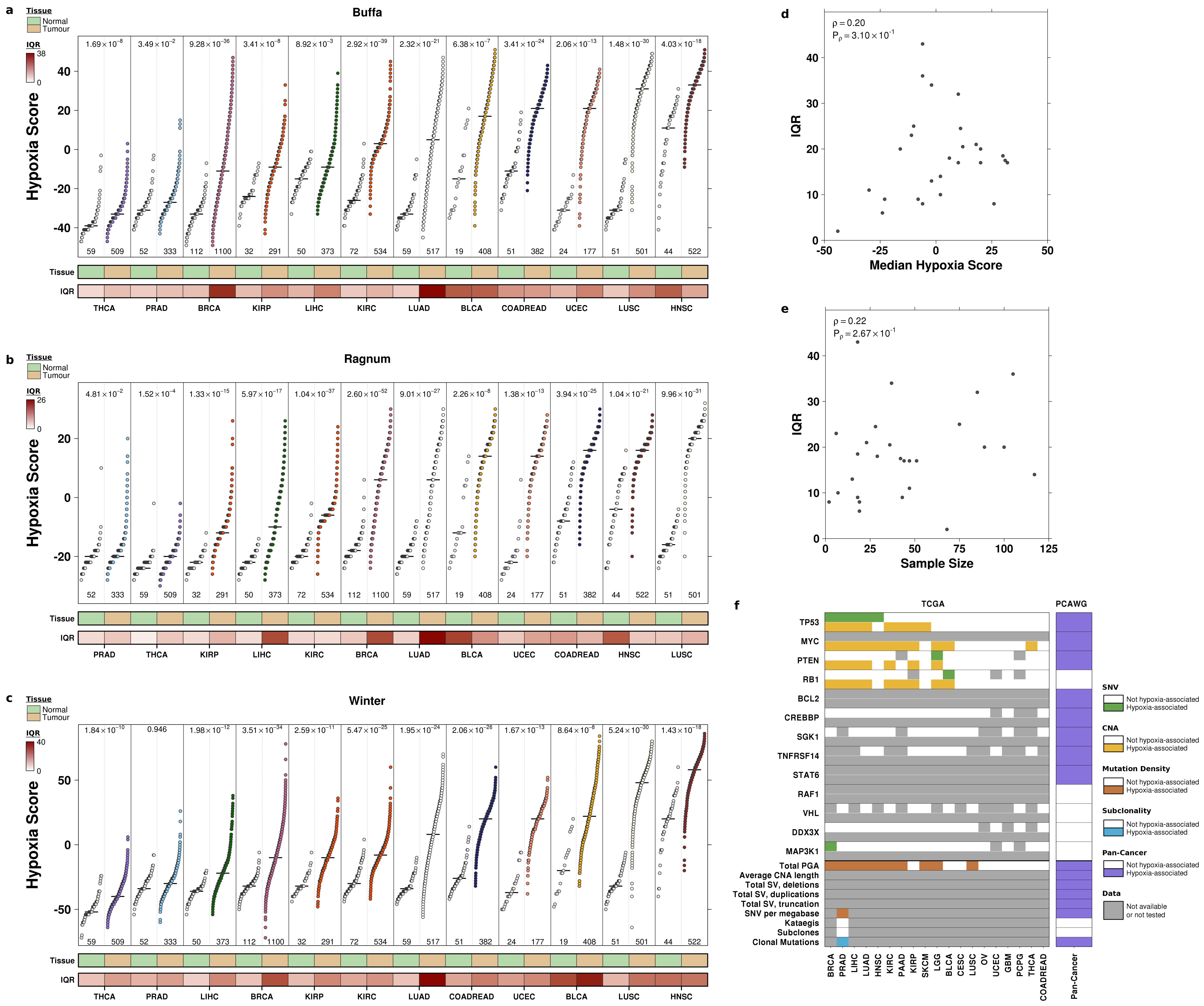
